# “Proteomic Insights into the Mechanism of Action of Maduramicin, a Novel Ionophore with Potent Antimalarial Activity”

**DOI:** 10.64898/2026.01.23.701272

**Authors:** Aakriti Singal, Hina Bharti, Santonu Kumar Pradhan, Deeptashree Nandi, Ravindra Varma Polisetty, Alo Nag

## Abstract

Malaria is a common infectious disease in tropical countries and poses serious health burden due to limited treatment choices. In recent years, many of the state of the art anti-malarials such as Artemisinin, Chloroquine and Primaquine have been rendered ineffective due to emergence of resistant parasites. This underscores the urgent need to develop new anti-malarials and elucidate their mechanism of action. Recently, we have demonstrated the potent anti-malarial properties of a known anti-coccidial ionophore, Maduramicin. In this study, we investigated the mechanisms of action of Maduramicin in *P. falciparum* by assessing its stage-specific activity and time-dependent effect on parasite development. The drug exhibited maximum anti-malarial activity against the schizont stage and was characterised as a slow-acting drug. To gain mechanistic insights, we employed iTRAQ based quantitative proteomics approach to analyse global proteome alterations in *P. falciparum* following Maduramicin treatment. Analysis of the differentially regulated proteins were carried out by using Gene Ontology and Kyoto Encyclopedia of Genes and Genomes pathway database. The data reflected significant perturbations in protein synthesis, energy metabolism, and other key metabolic pathways in response to Maduramicin treatment. Collectively, our results of the proteomics study were validated by quantitative RT-PCR analysis of 9 representative genes. Our findings provide the basis for understanding the lethal activity of Maduramicin on *P*.*falciparum*.

## INTRODUCTION

Malaria continues to be one of the most prevalent infectious diseases, affecting millions of lives across the globe. The protozoan parasite *Plasmodium falciparum* is the primary culprit, particularly affecting Sub-Saharan Africa and Asia (1). Extensively used anti-malarials such as Quinine, Chloroquine and Artemisinin have been rendered less effective with time due to escalating parasite resistance. Over the last two decades, Artemisinin based combination therapy (ACT), vector control, usage of insecticides and implementation of new vaccine regimen have reduced the malaria-related mortality by half (2,3). Although these approaches have shown promise, the emergence of the resistant form of *P. falciparum* has significantly challenged our progress with management of the disease and, therefore, incidences of malaria are again on the rise (4). This underscores the urgency for pioneering, alternative anti-malarial drug approaches. In recent years, carboxylic ionophores have been established as anti-plasmodial drugs (5). Their therapeutic potential has been reported in the veterinary field, particularly for poultry coccidiosis treatment (6,7), have demonstrated potential in combating malaria. Notable carboxylic ionophores such as Salinomycin, Nigericin and Monensin block the parasite transmission by hindering oocyst development and sporogony (8), thereby disrupting parasite transmission. Another ionophore, Maduramicin, a fermentation product of *Actinomadura yumaensis*, has garnered attention for its diverse therapeutic roles (9). Originally employed as an anti-coccidial agent, Maduramicin has displayed efficacy against cancer cells (10) and in Severe Combined Immune Deficient (SCID) (11). The drug has also been repurposed as an anti-malarial, showing effectiveness against various stages of the parasite’s lifecycle (12).

Maduramicin is a natural chemical compound initially isolated from the actinomycete *Actinomadura rubra*. It belongs to the class of polyethers, characterized by a series of electronegative crown ethers that possess a strong affinity for metal ions. The binding of these metal ions induces structural changes in the compound, exposing a hydrophobic methyl group. Its anti-parasitic mechanism centers on this exposed methyl group’s unique ability to form lipophilic complexes with various cations, particularly potassium (K+), sodium (Na+), and calcium (Ca2+). This facilitates the rapid translocation of these ions across cellular membranes (13, 14). Consequently, this disruption in ionic balance results in increased osmotic pressure, leading to dysregulated mitochondrial function and eventual demise of the parasite (15). When paired with other ionophore like compounds like pyrazoleamide PA21A050, Maduramicin enhances gametocytocidal activity (16) underscoring its potential in combination therapy.

Proteomics serves as a powerful tool for understanding the parasite response to therapeutics and its intricate interactions with the host (22). This methodology has facilitated the identification of proteins that function as specific biomarkers, potential targets for chemotherapy, and those that are differentially expressed in distinct parasite strains or under stress-induced conditions (23,24,25). Proteomic approach has helped vastly to assess the mechanism of action of anti-malarials such as chloroquine and artemisinin using gel-based analysis, MS analysis and isoleucine-based Stable Isotope Labelling SIL (26). Furthermore, the integration of isobaric tags for relative and absolute quantification (iTRAQ) with liquid chromatography-tandem mass spectrometry (LC–MS/MS) has facilitated a more comprehensive and profound exploration of the parasite’s proteome (68,69).

In this current study, we evaluated the impact of Maduramicin on distinct stages of the parasite’s life cycle and its overall progression. Furthermore, we have explored the alterations in the proteome of *P. falciparum* triggered by Maduramicin drug treatment, employing iTRAQ-coupled mass spectrometry techniques. The resultant proteomic dataset was meticulously examined to pinpoint proteins undergoing dysregulation, which were subsequently corroborated through quantitative real-time polymerase chain reaction (qRT-PCR) validation. Our findings provide novel mechanistic insights into the anti-malarial activity of Maduramicin.

## METHODS

### *In-vitro* culturing and synchronization of parasite

*P. falciparum 3D7* was cultured with human O^+^ human erythrocyte (5% hematocrit) in RPMI-1640 (Gibco, ThermoFisher Scientific, Waltham, MA, USA) medium supplemented with 0.5% (w/v) Albumax (Gibco, ThermoFisher Scientific, Waltham, MA, USA) in mixed gas environment (5% O_2_, 5% CO_2_ and 90% N_2_) at 37 °C (27, 67). Human whole blood was procured from White Cross Blood Bank, New Delhi. Under sterile conditions, erythrocytes were obtained by removal of plasma and peripheral blood mononuclear cells (PBMCs) using histopaque gradient. Parasitemia levels were routinely assessed using Giemsa staining of blood smears.

Synchronization was performed using D-sorbitol (Sigma-Aldrich, MO, USA) as described earlier (12, 67). Parasitized RBCs were collected and treated again with D-sorbitol, and the sorbitol was washed out using the culture medium. The resulting pRBC was stained and examined microscopically, and was found to be highly synchronized at the ring-stage (more than 80% of total PRBCs were in the ring-stage). Parasites of the ring-stage thus obtained were cultured in the medium for 16 h to obtain trophozoite, and 24 h for the schizont.

### Assessment of Stage Specific Activity of Maduramicin

Parasite was synchronized using Sorbitol lysis as mentioned above and the specific activity was performed as per published protocol mentioned previously with minor changes (66). The synchronized pRBC was exposed to different amounts of Maduramicin at 0.5% parasitemia and 2% haematocrit in the culture medium at 0h (ring stage), 16h (trophozoite stage) and 24h (schizont stage). Drug exposure studies were performed in a 96-well plate format over a concentration range 0.25-4.5ng/ml. Plates were kept in modular incubators for 24 h in mixed gas environment at 37°C. Thereafter, SYBR Green assay was performed with all the wells to calculate the percentage parasite inhibition. After 24 h, 100 μl of RBC lysis buffer (20 mM Tris base, 5mM EDTA, 0.0008% saponin, 0.08% Triton X-100, pH 7.5 with 0.2 μl SYBR® Green I/ml of lysis buffer) was added to each well. Plates were incubated in the dark for 1 h and then read for fluorescence (excitation 485 nm, emission 535 nm) on a TECAN Infinite F200 Pro plate reader. The data analysis was performed by MS Excel.

### Speed of action studies

The speed of action of the anti-malarial candidates was verified by the incubation of the synchronized parasites at the ring stage in the presence of Maduramicin at a concentration that was tenfold greater than the IC50 value. Parasites in thin blood smears were stained with Giemsa and observed under the microscope after 0, 8, 16, 24 and 36 h of incubation (30). .

### Protein Extraction from *Plasmodium falciparum*

Synchronized parasite at the schizont stage was exposed to 2.5ng/ml Maduramicin for 24h and protein were extracted from the parasitized RBCs. Control and treated groups consisted two biological replicates for the iTRAQ experiment. iRBCs were lysed by 0.1% saponin (Sigma) for 5 min. Free parasites were sedimented by centrifugation (8000 rpm for 5 min) and washed thrice with 1X PBS. Parasites were resuspended in lysis Buffer (100 mM Tris-HCl buffer (pH 7.4), 20mM NaCl, 70mM KCl, 12mM MgCl_2_, 1mM PMSF, 1mM DTT and 0.5%NP-40) and disrupted by ultrasonication (Misonix Ultrasonic Liquid Processer) for 5 min on ice at maximum amplitude. After centrifugation (12,000 rpm for 30min at 4°C), soluble protein fractions were recovered from the supernatant and protein content was estimated using Bradford reagent. 250µg of the fraction was precipitated using tri-chloroacetic acid to remove lipids and finally, the contents were completely dried using freeze drier.

### iTRAQ labelling and strong cation exchange

After protein precipitation in acetone, the pellet was dissolved in dissolution buffer. 100µg was taken in 20µl dissolution buffer, reduced, alkylated, trypsin-digested and labelled using the iTRAQ reagents four-plex kit according to the manufacturer’s instructions (Applied Biosystems). The resulting peptide solutions from control were labelled with iTRAQ 114 and 115 tags and Maduramicin-treated soluble protein samples were labelled with iTRAQ116 and 117 tags, and incubated at room temperature for 1 h. Labelled peptides were then pooled and acidified by mixing with the cation buffer load iTRAQ reagent for a total volume of 1 ml. The peptide mixture was subsequently fractionated by strong cation exchange (SCX) chromatography. The elution was monitored by absorbance at 214 and 280 nm, and 7 fractions were collected. Fractions were further analyzed by chromatography analytical column with specification ChromeXP, 3C18-CL-120, 3 μm, 120 Å and 0.3 × 150 mm. The used flow rate was 5 uL/min for the analytical chromatography to separate the peptides in the continuous gradient of elution with the 2-90% acetonitrile for 87 min total run time. The system used the solvent composition with a mixture of two reservoirs. Reservoir A of 98% water and 2% ACN in 0.1% FA, and B with 98% ACN and 2% water in 0.1% FA was used. The acquisition was executed with conventional data-dependent mode while operating the instrument name into an automatic MS and MS/MS fashion. The parent spectra were acquired with the scan range of 350-1250 m/z. Product acquired with the scan range of 100-1600 m/z. The ion source was operated with the following parameters: ISVF = 5500; GS1 = 25; GS2 = 22; CUR = 30.

### Identification of protein using iTRAQ based database

Protein quantification was carried out using the Protein Pilot software (Version 5.0, AB SCIEX) and a *P. falciaprum* database was utilized. Protein identification was performed with methylmethanethiosulfonate, selected for cysteine modification and iodoacetamide as a biological modification. The false discovery rate (FDR) analysis was calculated with an automated decoy database search strategy, embedded in the PSPEP (Proteomics System Performance Evaluation Pipeline) software that was integrated in the Protein Pilot software. The peptides in our iTRAQ labeling were then selected for protein quantification using unique peptides during the search. Only those peptides that had a global FDR of ≤1% were considered for subsequent analysis. The Protein Pilot software was exploited to generate the p values, based on the peptides used to quantify the respective proteins. Finally, the fold change (FC) was calculated as the average ratio of 116/114 and 117/115 for differentially expressed proteins and samples with a fold change of >1.2 or <0.8 and p < 0.05 were considered to be significantly differentially expressed proteins in control and Maduramicin-treated *P. falciparum* culture.

### Analysis of iTRAQ data using bioinformatics analysis

The NCBI GI (GeneInfo Identifier) numbers of identified *P. falciparum* proteins identified were converted into standard gene names for retrieval from the UniprotKB ID module. Then, in UniprotKB, PlasmoDB accession numbers of proteins were retrieved for further analysis. Biological processes, molecular function, and subcellular localization of differentially expressed proteins were assessed using Gene Ontology (GO) annotations downloaded from PlasmoDB. We also annotated protein pathway for significant proteins by searching Kyoto Encyclopedia of Genes and Genomes (KEGG) database. In addition, the protein-protein interaction networks of the significant proteins were analyzed by STRING (Search Tool for the Retrieval of Interacting Genes/Proteins) database (http://string.embl.de/).

### Quantitative real-time PCR

The parasite cultures used for the proteomic analysis were also used to perform quantitative RT-PCR (qPCR). Total RNA was extracted with TRIZOL® reagent following the manufacturer’s recommendations (Invitrogen, Waltham, Massachusetts) and treated with DNase (DNAfree Ambion). Total RNA was quantified using NanoDrop ND-1000 (Labtech). Total (1 µg) was reverse transcribed with the High-Capacity cDNA Archive Kit as described by the manufacturer (Applied Biosystems, Waltham, Massachusetts). The sets of primers (Sigma, Asao-ku, Kawasaki) used in this study are provided in the Supplymentary Table 2. Real time transcript quantification was performed using a QuantStudio 5 Real-Time PCR system (Applied Biosystems, Waltham, Massachusetts). Relative expression of the targets was determined using 2^-ΔΔCt^ method as previously described (32), 18s was used as internal control. All data were expressed as means ± standard deviation. A two-tailed Student’s t-test was employed to compare RT-PCR gene expression levels. Statistical significance was defined as p < 0.05.

### Western Blot Analysis

The parasite cultures used for the proteomic analysis were also used to perform western blotting. Following treatment, culture was pelleted and cell pellets were saponin lysed and washed thrice with ice-cold PBS. Subsequently, these parasite pellets were further lysed in radioimmunoprecipitation assay buffer (RIPA buffer) for 45-60 min and subjected to 8-10 rounds of sonication of 2s ON and 2s OFF. The lysates were centrifuged, the supernatant was collected and the protein content was determined by Bradford assay. Equal amounts of parasite lysates were loaded and resolved in SDS-PAGE and subjected to western blotting with Hsp70 (4876S, Cell Signaling Technology) and β-actin (sc-47778, Santa Cruz Biotechnology)) antibodies.

## RESULTS

### Effect of Maduramicin against blood stages of *P. falciparum*

In our previous research findings, we established that free Maduramicin exhibited a notable inhibitory effect on parasite progression, with an IC50 of 2.5 ng/ml (12). To further characterize its anti-plasmodial activity, we assessed its stage specific effects by treating synchronized ring, trophozoite and schizont stages of *P. falciparum* with varying concentrations for 24h. Remarkably, our investigations revealed that Maduramicin displayed the highest efficacy against the schizont stage of *P. falciparum*. Particularly, it demonstrated a 50% inhibitory effect at a concentration of 0.25 ng/ml when exposed to the later stages of Plasmodium development. Hence, Maduramicin exhibited the capability to target and eliminate all stages of *P. falciparum*, with the schizont stage showing heightened susceptibility to Maduramicin treatment compared to the early stages (see Figure 1).

**Figure 1.**
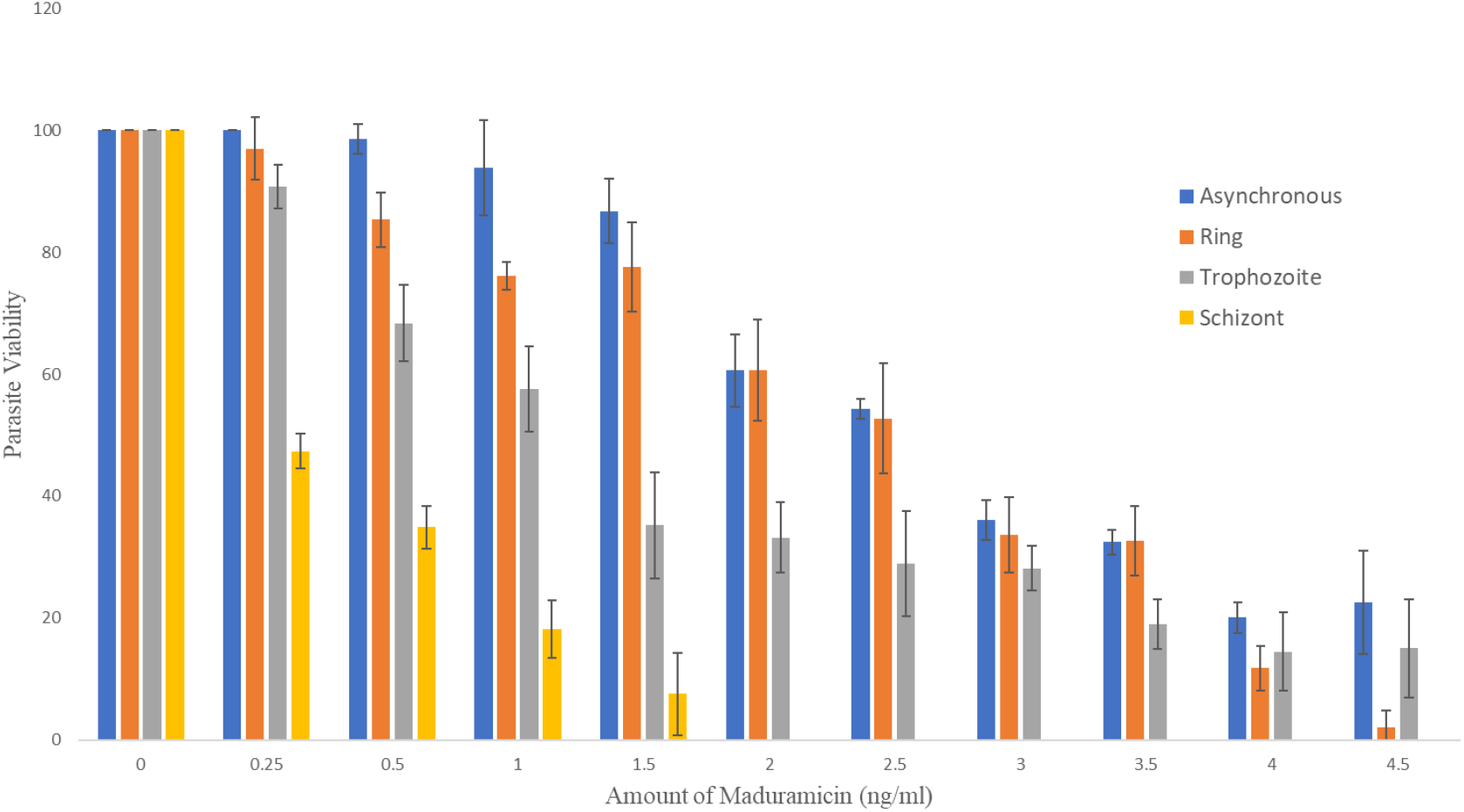
Stage specific activity of Maduramicin against asexual stages of P. falciparum 3D7. Synchronized (ring, trophozite and schizont) along with asynchronous parasite culture were exposed to different concentrations of Maduramicin for 24h. Graphical representation of the percent viability of asynchronous and synchronized (ring, trophozoite and schizont) parasite culture with increasing concentration of Maduramicin or vehicle control *** p < 0.05. Data are expressed as the means ± standard deviations of three independent experiments.

### Determination of speed of action of Maduramicin

To gain deeper insights into the anti-plasmodial attributes of Maduramicin, we conducted an evaluation of its speed of action against the asexual intraerythrocytic stages of *P. falciparum*. Our assessment encompassed an entire maturation cycle of the parasite, spanning from the ring to schizont stage, within the confines of the assay conditions. Notably, after 16 h of incubation, young trophozoites were observed. Strikingly, under the influence of Maduramicin, the parasite exhibited a distinct slowdown in its progression towards the schizont stage. Furthermore, the parasite’s ability to transition to the subsequent stage of the cycle was profoundly hindered in comparison to the control group (refer to Figure 2). A detailed morphological analysis of the parasite depicted a notable alteration – following drug treatment, the parasite was constrained to one end of the erythrocyte and its growth was markedly suppressed. These observations collectively underscore that Maduramicin exerts its impact predominantly at the schizont stage and qualifies as a slow-acting inhibitor of in vitro parasite growth.

**Figure 2.**
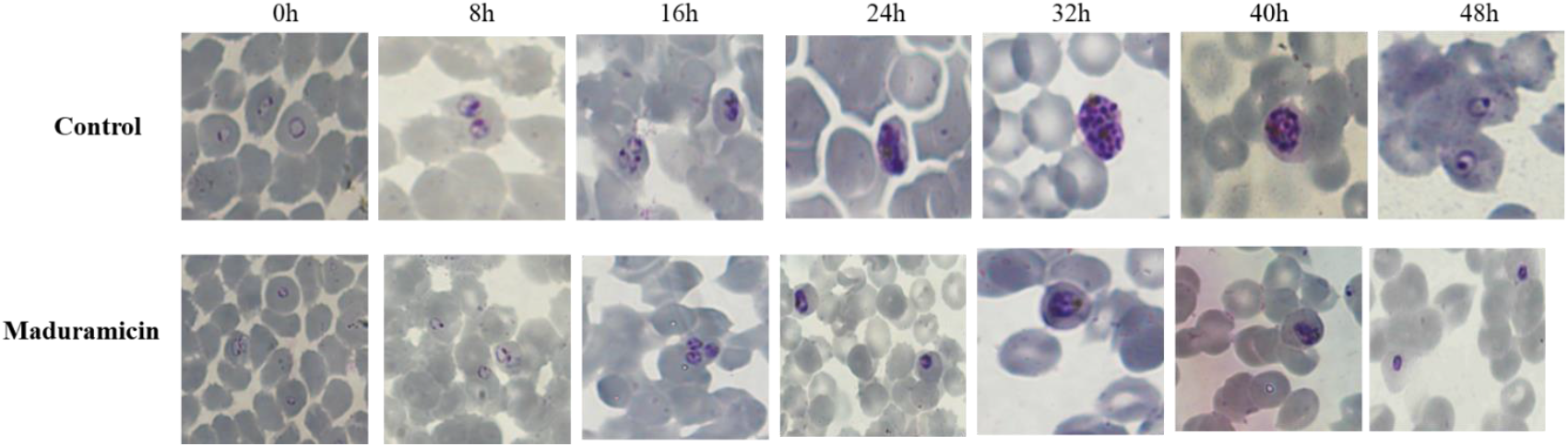
Speed of action investigation for Maduramicin. Synchronized parasites were exposed to either vehicle control or Maduramicin (ten times the IC50 concentration) and the effect on growth and morphology was evaluated for one complete maturation cycle [starting from ring (0 h), trophozoite (16–24 h) to schizont (40 h)]. Giemsa-stained blood smear micrographs are represented through images.

**Figure 3.**
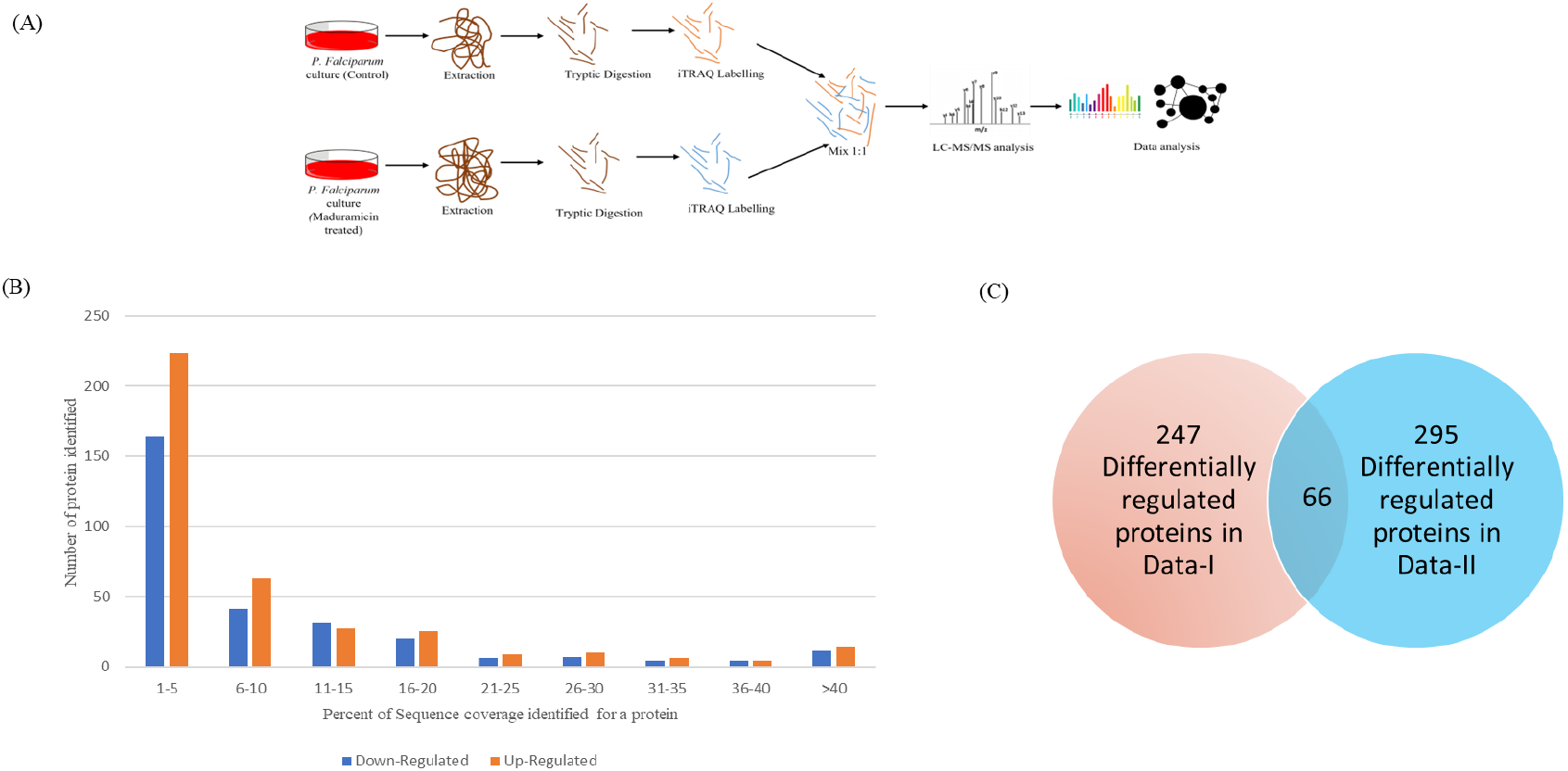
Information on the identified proteins. (A) Workflow for the experiment performed to identify the parasite proteins regulated by the parasite. The same protocol was followed for the biological replicate with different iTRAQ tags (B) The distribution of the identified proteins on the basis of sequence coverage. (C) A Venn diagram of the number of differentially regulated proteins identified in two biological replicates.

### Investigation of parasite response to Maduramicin treatment by iTRAQ

For identifying differential protein expression, proteomic approach coupled with LC-MS/MS was applied to Maduramicin treated *P. falciparum* culture. Two groups of schizont stage *P. falciparum* parasites were prepared: vehicle control and 2.5ng/ml Maduramicin (IC50)-treated groups. The proteins were extracted from each experimental group after 24 h of treatment. Peptides generated from trypsin digestion of control and Maduramicin treated *P. falciparum* lysates were labelled using the iTRAQ labels (control 114 and 116; Maduramicin treated; 115 and117), mixed together and analysed by liquid chromatography coupled to mass spectrometry (Figure 4). 963 proteins were recognized in both the samples. In order to explore the effect of Maduramicin on *P. falciparum* at the proteome level, differentially regulated proteins were screened according to the intensity of the iTRAQ reporter ions. From the biological replicates, 313 and 361 proteins were differentially regulated between the control and the treated groups of the parasite. Among the biological replicates, both the iTRAQ results depicted an overlap of 66 proteins, 37 down-regulated proteins and 29 up-regulated proteins on a basis of a fold change <0.8 (down-regulated) or >1.2 (up-regulated) respectively for the differentially expressed proteins (Table 2). The flowchart in figure 5 illustrates the funnelling of the high-throughput data obtained from proteomics.

**Table 1:** Data depicted from Gene Ontology annotation of the differentially regulated proteins.

| Gene ID | Name | Predicted Function | Biological Process | Localization |
| --- | --- | --- | --- | --- |
| PF3D7_0302900 | Exportin-1 | transporter activity | Regulation of protein transport<br>regulation of cellular localization<br>ribosomal subunit export from nucleus | cytoplasm<br>nucleus |
| PF3D7_1203700 | Nucleosome assembly protein | histone binding<br>chromatin binding | nucleosome assembly | nucleus |
| PF3D7_1136800 | DnaJ protein | Hsp70 protein binding | ubiquitin-dependent ERAD pathway<br>'de novo' protein folding<br>chaperone-mediated protein folding<br>cellular response to misfolded protein | endoplasmic reticulum<br>membrane<br>plasma membrane<br>vacuole |
| PF3D7_1464600 | Serine/threonine protein phosphatase<br>UIS2 | ferrous iron binding |  |  |
| PF3D7_0903700 | Tubulin alpha chain | anion binding<br>structural molecule activity<br>carbohydrate derivative binding<br>purine nucleotide binding | microtubule cytoskeleton organization<br>mitotic nuclear division | cytoplasm<br>microtubule |
| PF3D7_0212900 | Arginyl-tRNA--protein transferase | catalytic activity, acting on a protein<br>transferase activity, transferring acyl groups |  | cytoplasm |
| PF3D7_1325100 | Phosphoribosylpyrophosphate synthetase | transferase activity, transferring phosphorus-containing groups | purine nucleotide biosynthetic process<br>ribose phosphate biosynthetic process | cytoplasm<br>transferase complex, transferring phosphorus-containing groups |
| PF3D7_1441400 | FACT complex subunit SSRP1 | histone binding<br>nucleosome binding |  | transcription elongation factor complex<br>nuclear chromatin |
| PF3D7_1011800 | PRE-binding protein | mRNA binding | gene expression<br>regulation of gene expression | cytoplasm<br>nucleus |
| PF3D7_0617800 | Histone H2A | DNA binding | transcription, DNA-templated<br>chromatin silencing | nuclear chromatin |
| PF3D7_1369500 | Uncharacterized protein | RNA cap binding<br>mRNA binding | nuclear-transcribed mRNA catabolic process, nonsense-mediated decay<br>mRNA splicing, via spliceosome | nucleus<br>protein-containing complex |
| PF3D7_0818900 | Heat shock protein 70 | heat shock protein binding<br>ATP binding<br>ATPase activity<br>unfolded protein binding | de novo' protein folding<br>chaperone-mediated protein folding<br>cellular response to unfolded protein | cytoplasm |
| PF3D7_0501100.1 | Heat shock protein 40 | chaperone binding<br>unfolded protein binding | de novo' protein folding<br>chaperone-mediated protein folding | cytosol |
| PF3D7_1016400 | Serine/threonine protein kinase, FIKK family | protein serine/threonine kinase activity | protein phosphorylation<br>intracellular signal transduction | cytoplasm<br>nucleus |
| PF3D7_0707700 | E3 ubiquitin-protein ligase, putative | chaperone binding, ubiquitin protein ligase activity | protein quality control for misfolded or incompletely synthesized proteins | cytoplasm |
| PF3D7_0714000 | Histone H2B | DNA binding | nucleosome assembly |  |
| PF3D7_0619400 | Cell division cycle protein 48 homologue, putative | ATPase activity |  |  |
| PF3D7_1412300 | Nuclear transport factor 2, putative |  | nucleocytoplasmic transport | nuclear pore central transport channel<br>plasma membrane<br>vacuole |
| PF3D7_1015600 | Heat shock protein 60 |  | protein folding |  |
| PF3D7_1439800 | Vesicle-associated membrane protein, putative |  | membrane organization<br>organelle localization<br>endoplasmic reticulum organization | endoplasmic reticulum<br>membrane<br>plasma membrane<br>vacuole |
| PF3D7_0417300 | LETM1-like protein, putative |  | cellular metal ion homeostasis |  |
| PF3D7_1105400 | 40S ribosomal protein S4 | RNA binding<br>structural constituent of ribosome | translational elongation | cytosolic small ribosomal subunit |
| PF3D7_1451100 | Elongation factor 2 | translation regulator activity<br>RNA binding<br>GTPase activity<br>ribosome binding | translational elongation | ribonucleoprotein complex<br>cytosol |
| PF3D7_1110500 | Vacuolar protein sorting-associated protein 35 |  | retrograde transport, endosome to Golgi<br>intracellular protein transport | late endosome<br>membrane protein complex<br>plasma membrane<br>vacuole |

**Figure 4.**
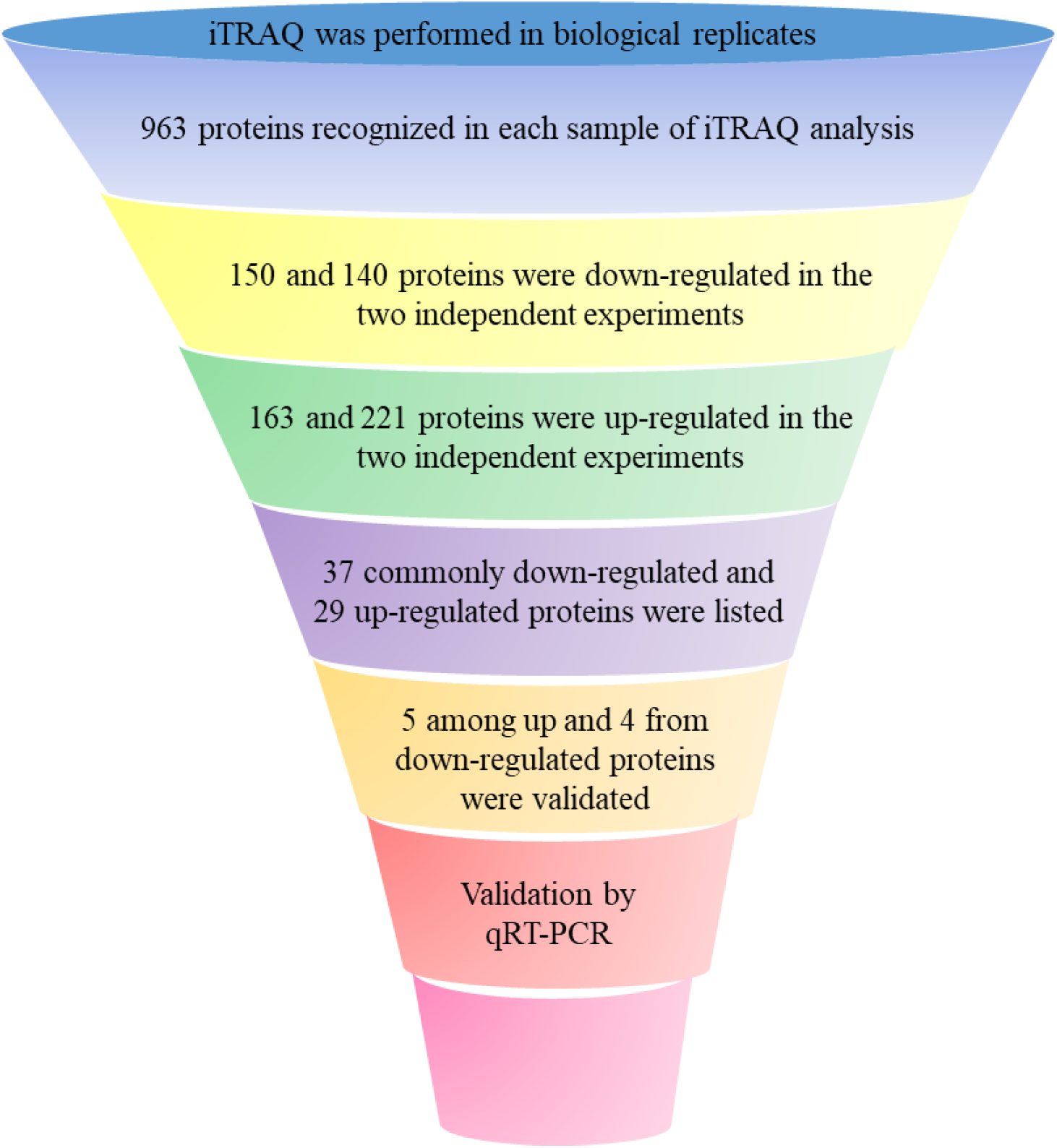
Flowchart for identification of differentially expressed proteins upon Maduramicin treatment of P. falciparum. A flowchart illustrating the series of deductions employed to select the annotated proteins that are most affected by Maduramicin.

**Figure 5:**
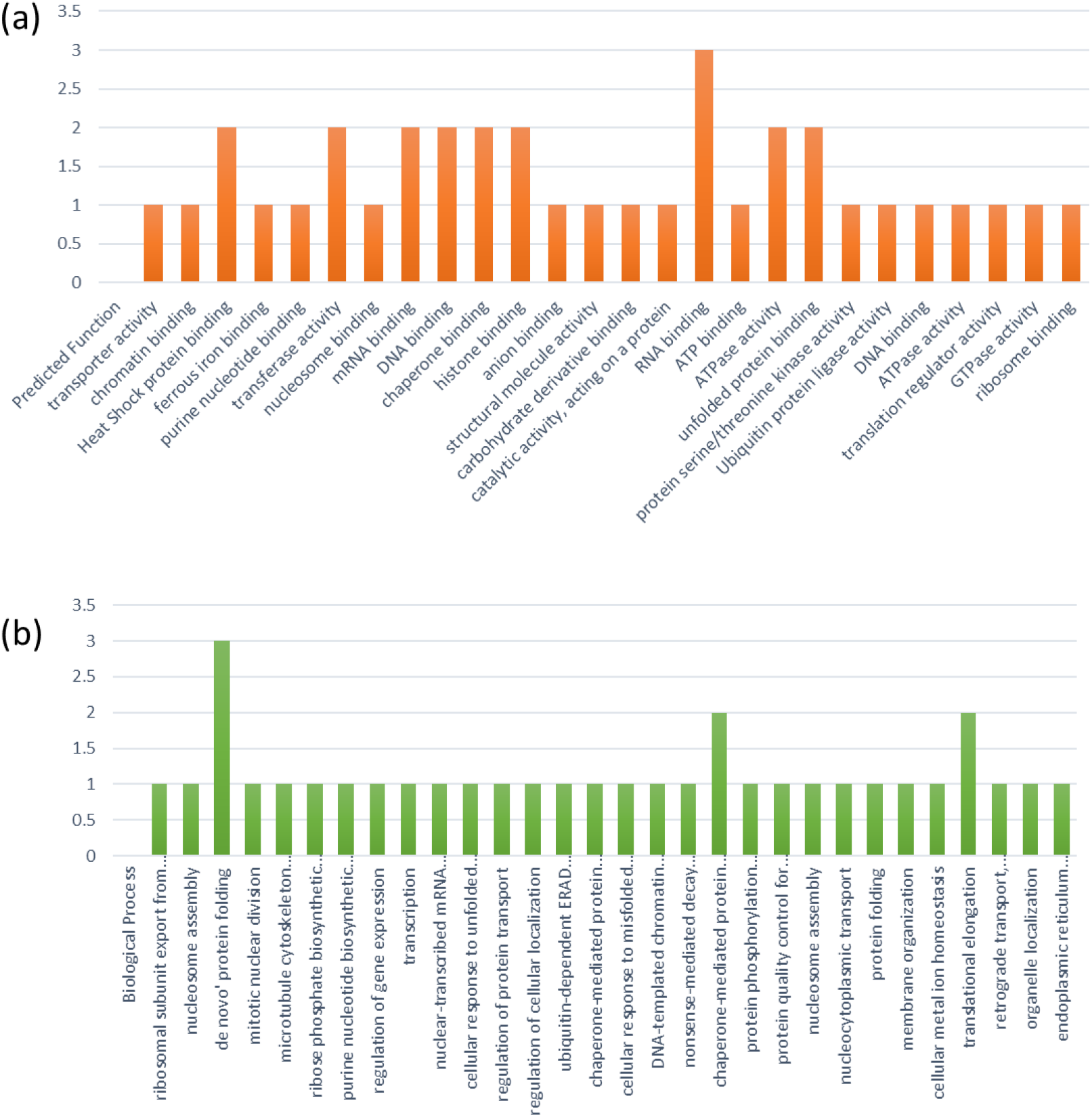
Number of differentially expressed proteins, identified by iTRAQ experiment, corresponding to different (a) functions and (b) biological processes by Gene Ontology. Write y-axis scale to whole numbers.

### Functional analysis of differentially regulated proteins via bioinformatic approaches

The differentially expressed proteins were annotated according to GO. Table 3 enlists the Gene ID and GO analysis of 27 differentially regulated proteins. GO analysis showed that the proteins can be classified into multiple categories of biological process, molecular function and cellular components. In the biological process analysis, maximum proteins are involved in de novo protein and chaperone mediated protein folding, followed by translation elongation and Hsp protein binding (Figure 6a). The proteins DnaJ, and Hsp60 involved in de novo protein folding and Hsp40 and Hsp70 involved in chaperone mediated protein folding are mostly found to be down-regulated. For molecular functions, most proteins such as 40s ribosomal protein S4, elongation factor 2 (EF2), PRE binding protein were implicated in RNA binding, while other proteins were involved in transferase activity (Arginyl transferase tRNA-protein transferase), mRNA binding, DNA binding (Histone H2A), chaperone binding (E3 ubiquitin protein ligase, Hsp40), histone binding (nucleosome assembly protein), ATPase activity (cell division cycle protein 48 homologue) and protein phosphorylation (serine/threonine kinase and phosphatase) (Figure 6b). In the classification of cellular components, a big chunk of differentially regulated proteins were cytoplasmic components, followed by nucleus, vacuole, plasma membrane, microtubule, and endoplasmic reticulum membrane (Figure 7). A schematic diagram is depicted in (figure 8), of the differentially regulated proteins according to their localization as obtained from GO. The above data reveals that Maduramicin insinuates its antimalarial effect via cytoplasmic proteins of the parasite that are involved in protein processing/synthesis.

**Figure 6:**
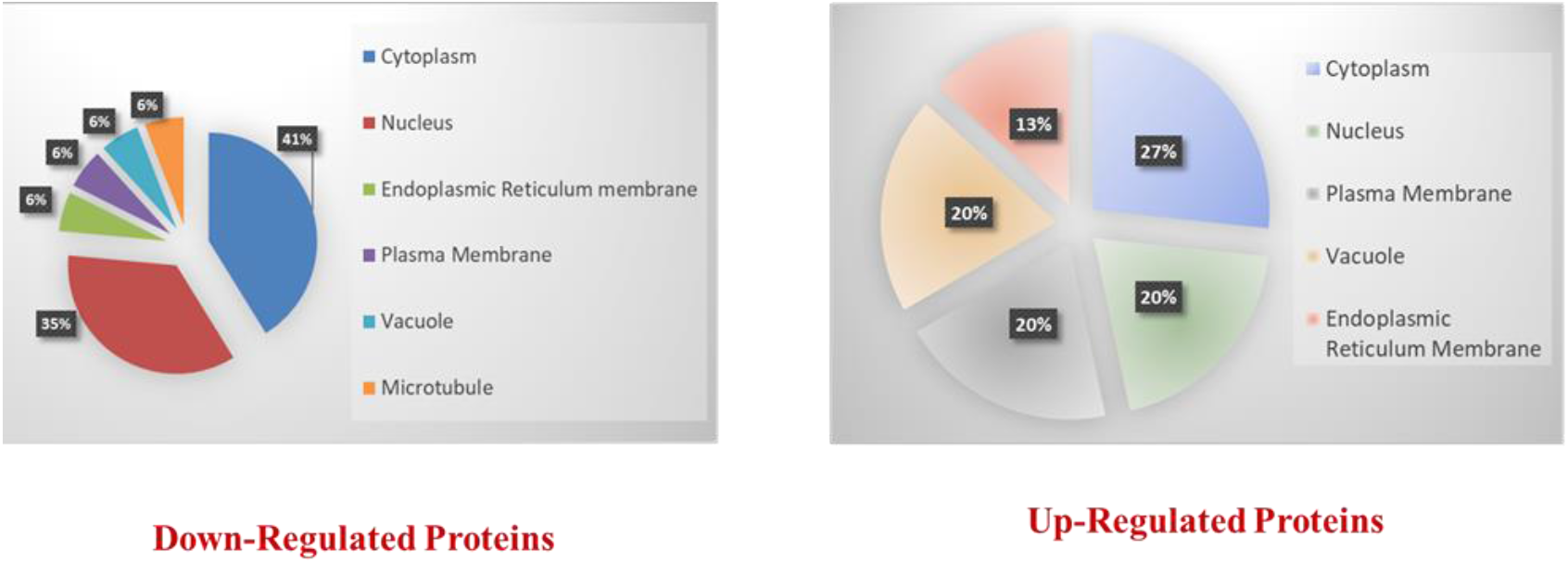
Classification on the basis of localization of differentially expressed proteins identified in Gene Ontology analysis of iTRAQ data.

**Figure 7:**
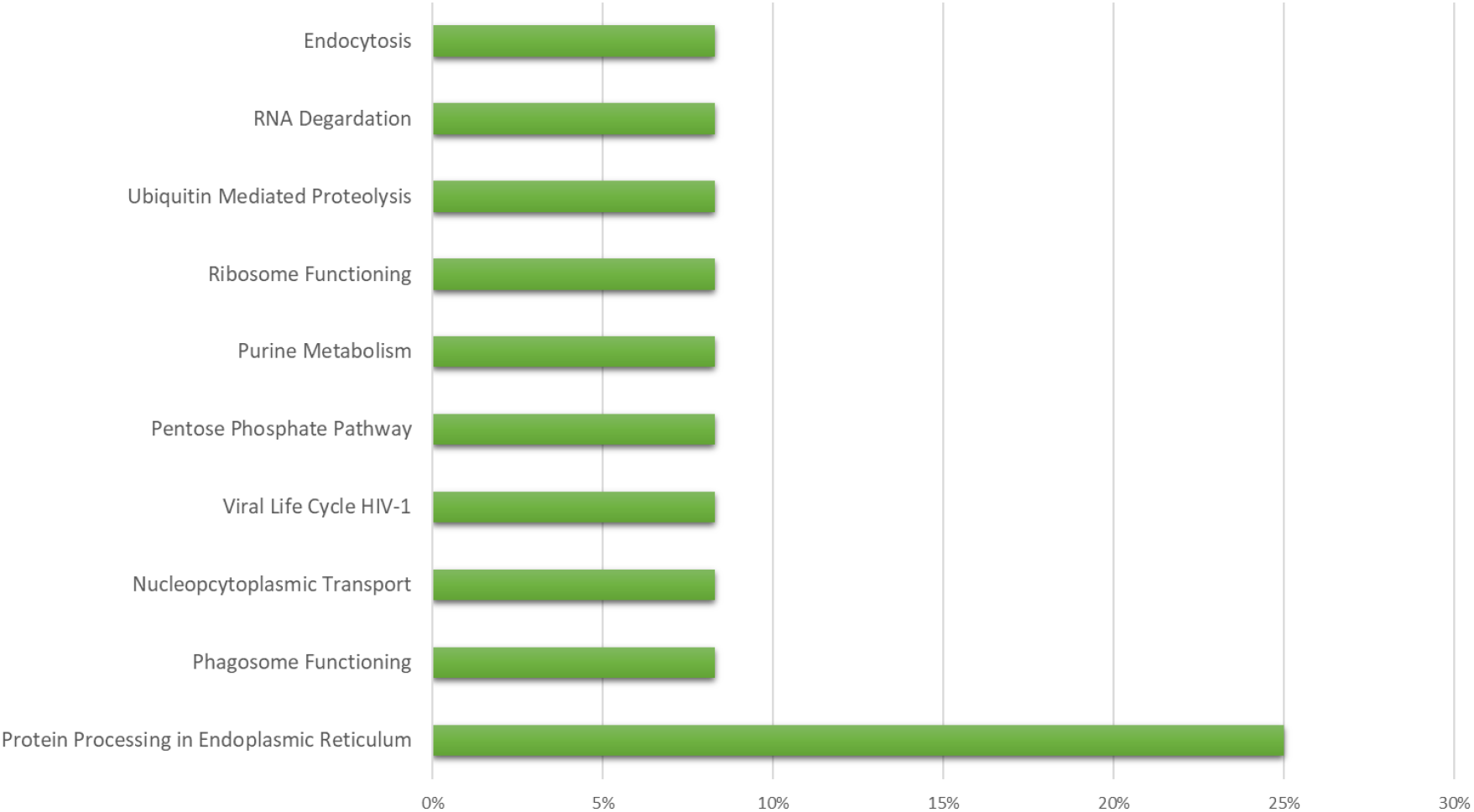
KEGG Pathway Analysis of differentially regulated proteins in P. falciparum under Maduramicin treatment.

**Figure 8:**
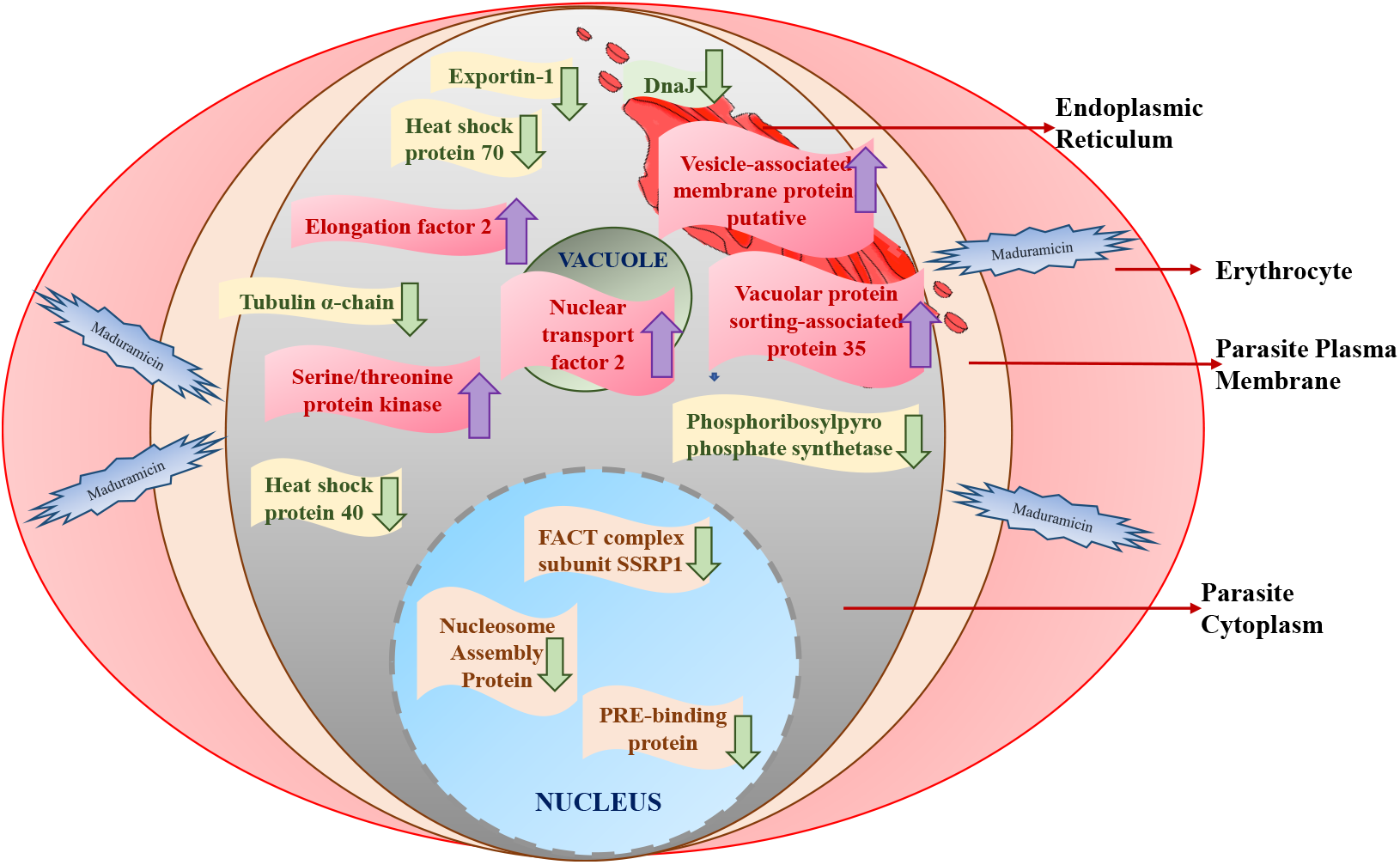
Overview of the proteins affected in P.falciparum upon Maduramicin according to their localization from Gene Ontology.

**Figure 8:**
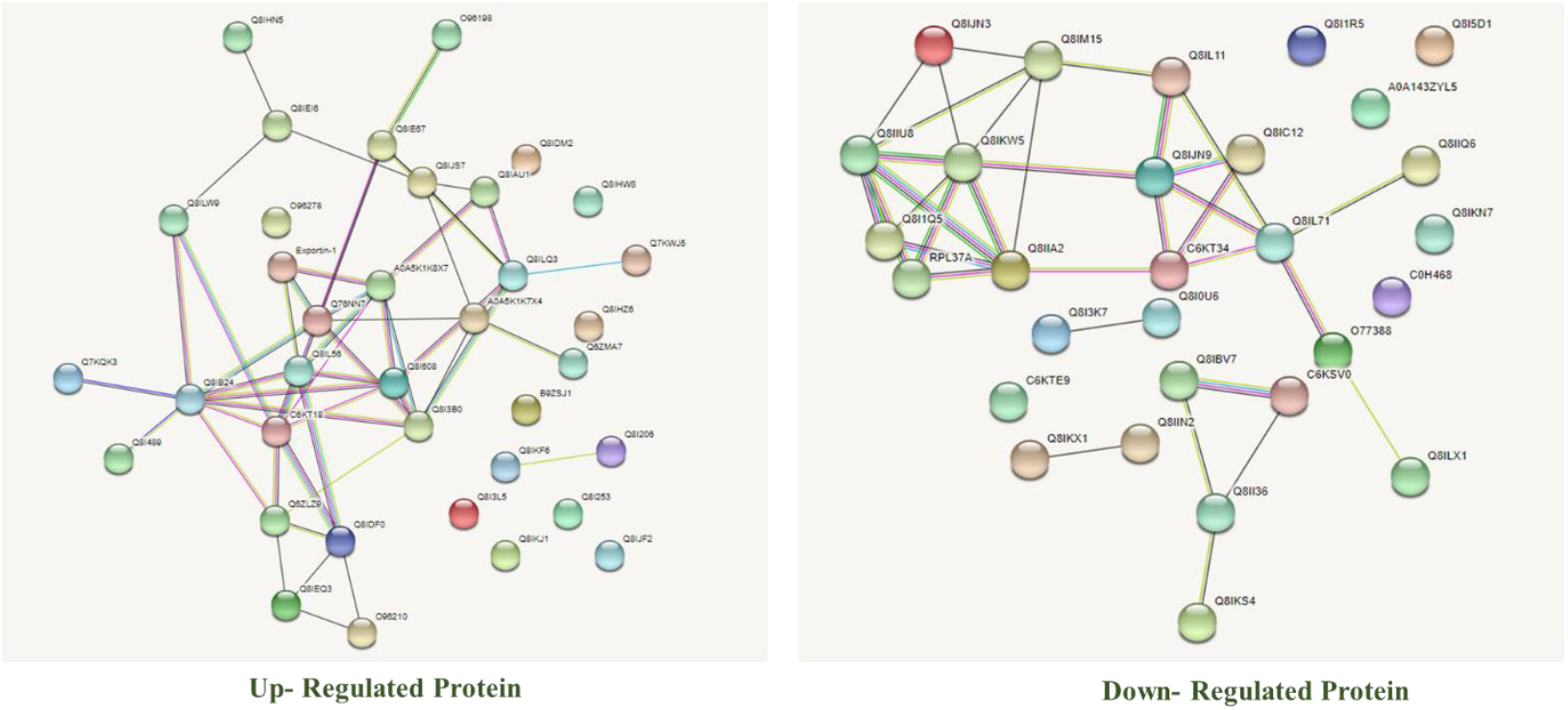
Protein–protein interaction (PPI) network of the differentially regulated proteins in the two biological replicates of parasite protein treated by Maduramicin in comparison to control group by STRING network. Lines in purple line color denote co-expression. Lines with orange color correspond to predicted interactions. Lines in blue color denote co-localization of the proteins. Yellow lines represent interaction based on shared protein domain of the proteins.

KEGG, a potent bioinformatics tool, plays a pivotal role in scrutinizing pathways in various scientific domains including systems biology, translational research, and omics investigations (65). The gene IDs corresponding to differentially regulated proteins have unveiled their participation in a rich tapestry of 22 functional pathways within KEGG (as illustrated in Figure 9). A substantial 14% of these proteins are notably enriched in the protein processing pathway in the endoplasmic reticulum, emerging as a focal point of perturbation induced by Maduramicin. Following suit are the metabolic pathways (10.7%), endocytosis (7.14%), RNA degradation (3.57%), ubiquitin-mediated proteolysis (3.57%), ribosome functionality (3.57%), purine metabolism (3.57%), pentose phosphate pathway (3.57%), viral life cycle HIV-1 (3.57%), nucleocytoplasmic transport (3.57%), and phagosome activity (3.57%).

**Figure 9:**
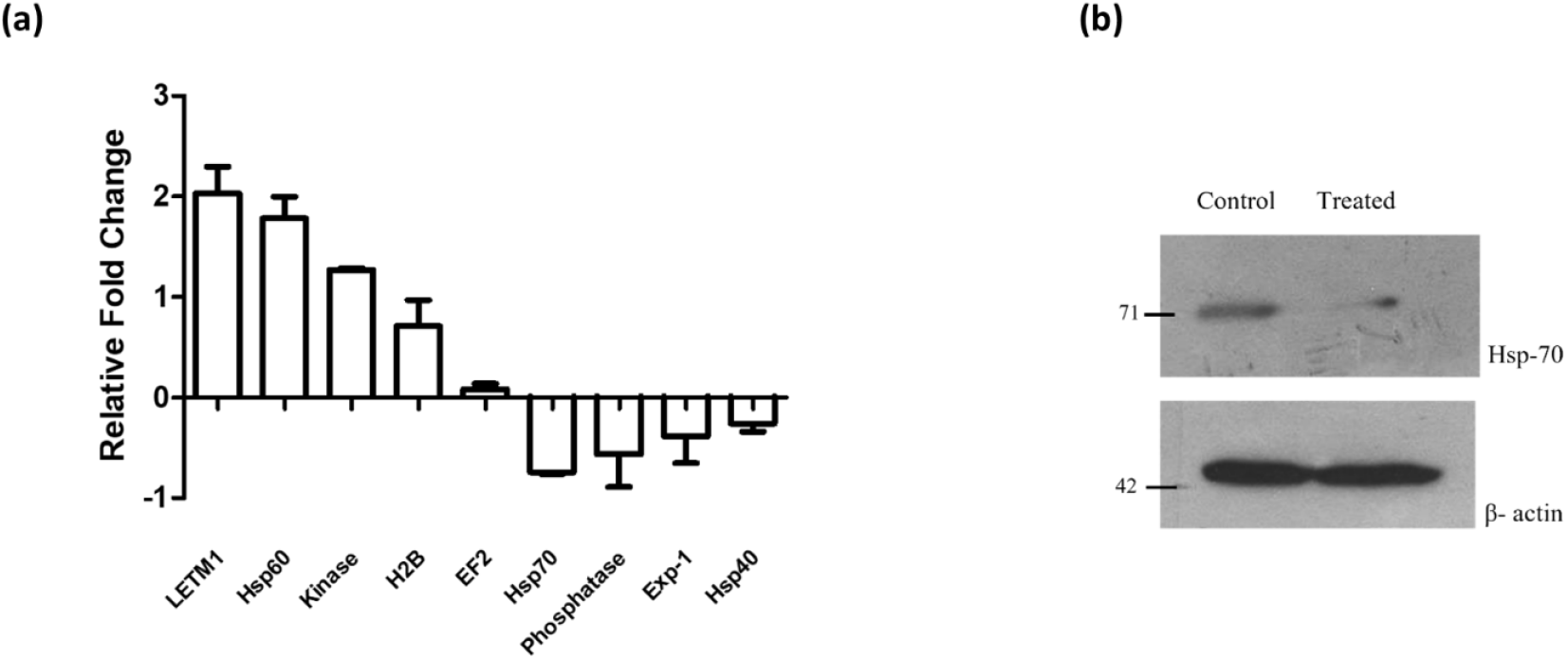
verification of iTRAQ Data. (a) Real Time PCR analysis of the levels of differentially regulated proteins under Maduramicin treatment in P. falciparum. The fold mRNA change was represented graphically. *** p < 0.05. Data are expressed as the means ± standard deviations of three independent experiments. (b) Western blot demonstrating inhibition of Hsp70 under Maduramicin treatment in P. falciparum.

The convergence of insights yielded by both GO and KEGG analyses forms an intricate and coherent tapestry. It resonates around the primary targets of Maduramicin, discernibly encompassing proteins implicated in nucleic acid binding, energy metabolism, and the facilitation of proper folding and stability for nascent proteins.

### Interaction network analysis of the differentially regulated proteins by Maduramicin

To unravel the intricate web of interactions among these proteins subject to differential regulation, we harnessed the power of STRING analysis (depicted in Figure 10). In the wake of Maduramicin treatment, a dynamic interplay emerges among parasite proteins, collectively weaving a comprehensive network that extends its functional influence across diverse biological processes, a remarkable adaptation to the prevailing stress conditions. Lines between nodes in STRING analysis represent various types of evidence for the association, such as neighborhood, gene fusion, co-occurrence, co-expression and protein homology. There is apparent interconnected network of 25 nodes for upregulated proteins that were mostly part of protein folding machinery. For the downregulated set, multiple small clusters of proteins with functions in various processes, including translation elongation and DNA-binding were present. Among the cohort of down-regulated proteins, Exportin-1 of *P. falciparum*, an integral player in intracellular transport, forms intriguing connections within the STRING network. This network reveals its indirect influence on Early Transcribed Membrane Protein 5, Peptidyl-prolyl cis-trans isomerase, and the FACT complex subunit SSRP1. This suggests that the inhibition of Exportin-1 intricately hampers the movement of several other critical proteins within the parasite cell.

Another pivotal protein in the network is Heat Shock Protein 70, which acts as a central hub, interlinking with Heat Shock Protein 40, Heat Shock Protein DNAJ homologue, actin-2, Peptidyl-prolyl cis-trans isomerase, FACT complex subunit SSRP1, Nucleosome Assembly Protein, 60s ribosomal protein L32, Histone H2A, and Tubulin alpha chain. This intricate interplay highlights the extensive impact of Heat Shock Protein 70 modulation on diverse cellular components and processes.

In the network of down-regulated proteins, two prominent nodes, 40S ribosomal protein S4 and EF2, emerge as interaction hubs. These nodes extend their connections to Serine/Threonine Protein Kinase, Eukaryotic Translation Initiation Factor 3 subunit M, 40S ribosomal protein S18, and Heat Shock Protein 60, implying their pivotal roles in orchestrating protein synthesis and cellular responses.

Conversely, the network of upregulated proteins demonstrates a higher degree of cross-reactivity. Notably, Heat Shock Protein 60 takes center stage once again, acting as a nexus that interfaces with E3 ubiquitin protein ligase (putative), vesicle-associated membrane protein, cell division protein 48 homologue, and M17 leucyl aminopeptidase. This intricate web of interactions suggests a coordinated response orchestrated by these upregulated proteins, potentially contributing to adaptive cellular changes under the influence of Maduramicin.

### Verification of the differentially regulated proteins by qPCR and western blotting

To validate the iTRAQ results, the transcriptional levels of 12 candidate differentially regulated proteins were analysed using qRT-PCR (Figure 11a and 11b). Among the proteins exhibiting the most notable fold changes, exportin-1 and histone H2B stood out and warranted further investigation. Given the significant impact of Maduramicin on heat shock protein expression according to proteomic data, we directed our attention towards heat shock proteins 40, 70, and 60 for validation. The intriguing reciprocal relationship between serine/threonine protein phosphatases and kinases, as observed in both proteomic and iTRAQ datasets, motivated us to validate the same.

Expanding our validation efforts, we included additional intriguing candidates such as DNA helicase, hexose transporter, hemolysin III, EF2, and LETM1-like protein. Through our verification experiments, we discerned a consistent trend: LETM1, Hemolysin III, Hsp60, Kinase, H2B, and EF2 exhibited upregulated expression, while DNA Helicase, hexose transporter, Hsp70, exportin-1, and Hsp40 demonstrated down-regulation. Importantly, these qRT-PCR findings harmoniously aligned with the initial iTRAQ results. Moreover, our western results (Figure 11c and 11d) indicated reduction in Hsp70 expression in Maduramicin-treated samples. This finding further supports the notion that Maduramicin exerts its action by impeding Hsp70, a chaperone protein pivotal for the function of its co-chaperone, Hsp40, which in turn plays a crucial role in parasite survival.

## DISCUSSION

The global fight to control malaria requires a multifaceted approach, which will be enabled by a better understanding of the biology of *P. falciparum (60)*. In 20^th^ century, Artemisinin alone or in combination with other drugs, has been central to anti-malarial therapy (34). Recently, ionophores have been increasingly used in Artemisinin Combination Therapy (ACT) (35). Maduramicin, in particular, shows promise, especially in liposomal formulation which enhances efficacy and offers prophylactic activity (12). In view of the recent advancements in the anti-malarial role of Maduramicin, we thought of elucidating its mechanism of action in *Plasmodium*. Studying its molecular mechanism will help us to identify immunomodulatory, anti-inflammatory, and antioxidant effects of the drug, optimize its dosing, treatment timing and patient selection. Furthermore, the benefits as well as risk factor associated with the consumption of the drug can also be assessed.

To begin with, the stage specific activity analysis of *P. falciparum* against Maduramicin revealed that the drug is most effective at the schizont stage unlike Quinines that are multi stage drugs and Primaquine that acts on the liver stage of the parasite (29). Maduramicin’s distinctive feature lies in its gradual inhibition of *in vitro* parasite growth which complements the rapid action of Artemisinin, a widely use anti-malarial (36). Owing to its potent impact on the schizont stage and its slow mode of action, Maduramicin holds promising clinical implications. Artemisinin primarily targets the ring stage and to a lesser extent the trophozoite stage of the parasite (37). By acting at different stages and speeds, Maduramicin can serve as a valuable component of ACT providing a multifaceted attack on the parasite’s lifecycle, enhancing the antimalarial efficacy.

To explore Maduramicin’s impact on the intricate cellular processes, we performed iTRAQ analysis revealing differential expression of 66 proteins, (37 downregulated and 29 upregulated). Known for inducing mitochondrial dysfunction, cytoplasmic vacuolization and affecting protein translation (61,62), Maduramicin also disrupts cytokine signaling, apoptosis, calcium pathways in other systems (63) and shows potential as an anti-cancer agent (10). Here, we found that Maduramicin treatment in *P. falciparum in-vitro* culture disrupted the energy metabolism and protein synthesis machinery of the parasite, akin to the proteomics result of Chloroquine treatment in *P. falciparum* (64).

GO and KEGG analysis indicated that Maduramicin affected proteins are enriched in pathways related to protein processing, RNA binding, chaperone mediated folding, nucleocytoplasmic transport, amino acid tRNA synthesis, DNA replication, and pentose phosphate pathway corroborate with the predicted functions obtained from GO such as, transporter activity, transferase activity, DNA binding and ATPase activity respectively. Maduramicin upregulated serine/threonine kinases and downregulated phosphatases, suggesting induction of parasite phosphorylation as a stress response (38). Since phosphatases are critical for DNA replication, schizont formation and RBC egress (39,40), their inhibition likely hampers parasite proliferation that contributes to parasite death.

Cellular response to stressful conditions majorly involves overexpression of Hsp proteins that facilitates correct folding of proteins (41). Similarly in *P. falciparum*, Maduramicin markedly altered Hsp 40 and Hsp70 levels, a chaperone-cochaperone complex essential for protein translocation, folding, assembly and parasite viability (46,47). Hsp 70 also regulates protein protein homeostasis via folding, disaggregation and ubiquitin-proteasome system (42,43), making it a potential therapeutic target, as its disruption impairs cell survival (44,45).

Another important protein, Exportin-1 was identified as a hub in the STRING network, was downregulated by Maduramicin treatment. It was found to be upregulated in the protein profiling of mefloquine resistant *P*.*falciparum* as compared to control. Exportin-1 is known to mediate nucleocytoplasmic export influencing drug resistance, cell cycle and transcription of the parasite (50,51). This novel finding reveals a potential mechanism by which Maduramicin exerts its anti-malarial effects by disrupting essential protein transport and impairing parasite survival and highlighting the complexity of the drug-parasite interaction.

Key proteins including DNA helicase, Hemolysin III and Hexose transporter were also validated as Maduramicin targets correlating with anti-parasitic effects. Helicases are highly versatile ubiquitous enzymes that are responsible for multiple essential cellular activities. Helicases are designated as the lifeline of cells since it participates in replication, translation, DNA repair, recombination and nucleic acid metabolism impacting the processes of proliferation, differentiation and embryogenesis. Therefore, helicases are vital for cell survival and can serve as potent drug targets. The observed reduction in helicase levels in *P. falciparum* upon Maduramicin treatment implies its role in hampering parasite replication. Agents inducing DNA-protein cross-links are known to stall helicases and polymerases, leading to replication blockage and helicases instability. Maduramicin may exert similar replicative roadblock, in the parasite. Subsequently, hexose transporters mediate the facilitated diffusion of glucose across the parasite plasma membrane that keeps the parasite active. Glucose deprivation disrupts progression through blood stages the parasite since it fails to meet the high energy demands of the early trophozoite stage. Compound 3361 that inhibits the hexose transporter has been recognised as an effective anti-malarial in almost all *Plasmodium* strains including *P*.*vivax, P. knowlesi, P. yoelii and P. falciparum*. Inhibition or low levels of Hexose transporter can cause glucose deprivation jeopardising the energy metabolism that can lead to killing of the parasite. Another upregulated target gene, **hemolysin** enzyme is capable of forming pores in the erythrocytes that leads to rupturing of the host cell organelles for the parasite to survive within the cell. It probably acts via channel formation (pore forming), detergent action, or lipase activity against the erythrocyte membrane. Therefore, upregulation of Hemolysin by Maduramicin probably causes hemolysis of the infected erythrocytes creating a hostile environment for the parasite. Considering the importance of target genes for the proper functioning of the metabolic system of an organism we propose the following model (figure 12). Proteomic studies of anti-malarials show that Chloroquine and Artemisinin upregulate oxidative stress proteins while Doxycycline affects glycolysis but none impact heat shock or ribosomal proteins (52,55,56), likely due to methodological differences. Verification of the iTRAQ analysis unveiled that Hsp 40, Hsp70, Serine threonine phosphatase, DNA helicase and glucose transporter and exportin-1 are found to be down regulated whereas LETM1, Hemolysin III, EF2, Hsp60, Serine threonine kinase and H2B are upregulated. As anticipated our qPCR data verification study aligns with the iTRAQ results. However, since the technique assesses the transcription level in the parasite, further in-depth biochemical studies are required at the protein level to confirm our premise.

**Figure 12:**
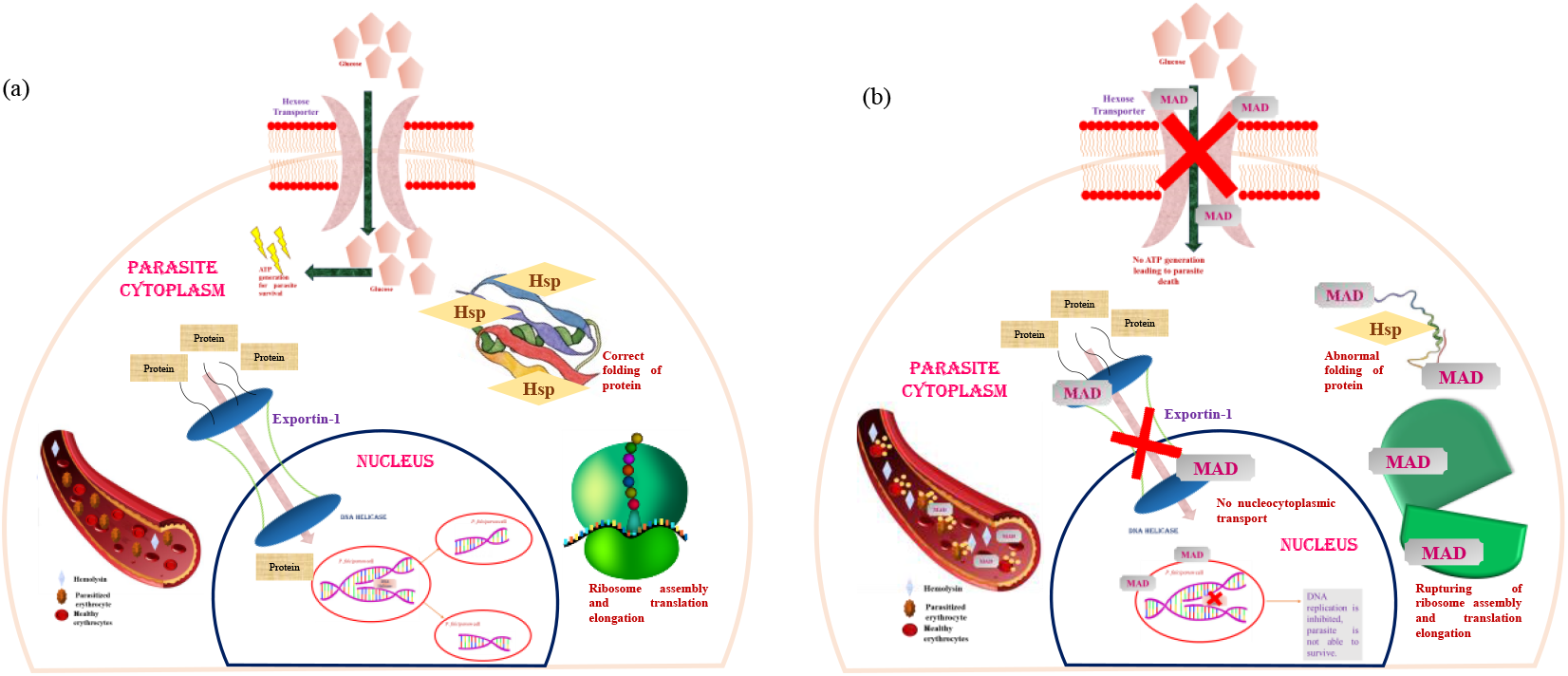
Schematic representation of Maduramicin-mediated antimalarial effects through various target proteins. (a) P. falciparum modulates the host proteome for its own survival. (b) Maduramicin (MAD) treatment of malarial cells perturbs the expression of the target genes thus hampering P. falciparum metabolism, ultimately culminating to the parasite death.

In summary, our investigation highlights the distinctive antimalarial properties of Maduramicin. It’s slow-acting nature and strong efficacy against the schizont stage distinguish it from other available antimalarials. Using iTRAQ-based quantitative proteomics, we provide first comprehensive insights into proteomic alterations in *P. falciparum* following Maduramicin exposue. Our study established that Maduramicin’s antimalarial activity is driven by disruption of key cellular processes including protein folding, DNA replication, energy metabolism, cell signalling, ribosome function, and nuclear-cytoplasmic transport within the parasite. Notably, upregulation of hemolysin III suggests membrane destabilization and hemolytic stress. Collectively, these perturbations disrupt protein homeostasis, metabolic regulation, and nucleocytoplasmic trafficking, culminating in growth arrest and parasite death. This multi-targeted, slow-acting mechanism highlights Maduramicin’s potential as a complementary agent to fast-acting antimalarials like artemisinin. By simultaneously targeting multiple vital pathways, Maduramicin emerges as a promising antimalarial candidate, offering valuable insights to guide future drug development and therapeutic interventions against malaria.

## Authors Contribution

AS designed and performed most of the experiments, critically analysed the results and prepared the manuscript. HB helped with some experiments and critical analysis of the results and manuscript review. SKP participated in data analysis, manuscript review and editing and DN contributed towards data interpretation and critical review of the manuscript. RVP supervised the proteomics related experiments, provided expert suggestions and reviewed the manuscript. AN contributed to overall supervision of the project, conceptualization, methodology, resources, critical data analysis, writing, review and editing the manuscript and fund acquisition. All the authors analysed the results and approved the final version of the manuscript.

