## Supplementary data for "“Proteomic Insights into the Mechanism of Action of Maduramicin, a Novel Ionophore with Potent Antimalarial Activity”"

This SI PDF file includes:

Tables S1 and S2

**Table S1:** Nucleotide sequence of primers used for qRT-PCR validation study.

| S.No. | Name of the Gene | Forward Primer | Reverse Primer |
| --- | --- | --- | --- |
| 1. | Exportin-1 | ATCACTAAAGGAATTCGCAA | TTATTCAACGTTTATACATTCGTC |
| 2. | Heat Shock Protein 70 | CCAACCAGGTGTCTTAATTC | GTTTACCAGTGGATTTTTCTA |
| 3. | Heat Shock Protein 40 | GTAATTGGTTCTAATTATGATAA | TACAGCATAGTAATCCGTTT |
| 4. | Serine/threonine protein phosphatase | CACATAGCTCAGATGAAAAT | CTTTTCTACATAAAATTTTATTCTAT |
| 5. | Serine/threonine protein kinase | TTTTTTTTTGTCTATATTTGATAAGG | GTTTTTTCATTTTTATAAGTATCTTC |
| 6. | Elongation Factor2 | TAGTTGAACCAATCTATCTTGTCGAT | ACCAGATGTAGCAGCACGTAAA |
| 7. | Heat Shock Protein 60 | TGCTATAAAAATGAATACATCAGATT | TACCACCACCTGGAACAATA |

**Table S2**. List of differentially regulated proteins by Maduramicin in P. falciparum commonly found in the two data sets.

| Gene Symbol | Name of the protein | Fold Change±S.D. |
| --- | --- | --- |
| HT1 | Hexose transporter | 0.34185±0.2 |
| PF3D7_0406800 | Sexual stage-specific protein | 0.3665±0.01 |
| FKBP35 | Peptidyl-prolyl cis-trans isomerase | 0.53725±0.24 |
| - | Uncharacterized protein | 0.4991±0.07 |
| XPO1 | Exportin-1, putative | 0.58405±0.16 |
| ACT2 | Actin II | 0.62565±0.21 |
| DNAH | DNA helicase | 0.5727±0.13 |
| PF3D7_0919000 | Nucleosome assembly protein | 0.6382±0.16 |
| DNAJ | DnaJ protein, putative | 0.58815±0.07 |
| ZFP | Zinc finger protein, putative | 0.6601±0.15 |
| PF3D7_1358900 | Serine/threonine protein phosphatase UIS2, putative | 0.6688±0.15 |
| ETRAMP5 | Early transcribed membrane protein 5 | 0.4622±0.17 |
| - | Uncharacterized protein | 0.603±0.03 |
| - | Uncharacterized protein | 0.62645±0.06 |
| PHIL1 | Photosensitized INA-labeled protein PHIL1 | 0.6841±0.12 |
| Tuba | Tubulin alpha chain | 0.6585±0.05 |
| K13 | Kelch domain-containing protein, putative | 0.68785±0.08 |
| DBP1 | ATP-dependent RNA helicase DBP1, putative | 0.6675±0.04 |
| Pfj4 | Heat shock protein DNAJ homologue Pfj4 | 0.674±0.05 |
| - | Uncharacterized protein | 0.68595±0.04 |
| RPL32 | 60S ribosomal protein L32 | 0.6699±0.004 |
| - | Uncharacterized protein | 0.6792±0 |
| ECT | Ethanolamine-phosphate cytidylyltransferase | 0.6824±0.013 |
| ATE1 | Arginyl-tRNA--protein transferase | 0.73225±0.06 |
| - | Uncharacterized protein | 0.7113±0.018 |
| PRPS | Phosphoribosylpyrophosphate synthetase | 0.7047±0 |
| SSRP1 | FACT complex subunit SSRP1 | 0.7047±0.01 |
| RBP | PRE-binding protein | 0.75545±0.034 |
| H2A | Histone H2A | 0.70515±0.037 |
| NARS | Asparagine--tRNA ligase | 0.72145±0.033 |
| PTP7 | EMP1-trafficking protein | 0.7657±0.02 |
| - | Uncharacterized protein | 0.7036±0.08 |
| HSP70 | Heat shock protein 70 | 0.6227±0.22 |
| PRESAN | PRESAN domain-containing protein | 0.7269±0.085 |
| - | Uncharacterized protein | 0.75275±0.06 |
| HSP40 | Heat shock protein 40, type II | 0.8017±0.01 |
| - | Uncharacterized protein | 1.6787±0.33 |
| - | Uncharacterized protein | 1.68655±0.36 |
| IMC1g | Inner membrane complex protein 1g, putative | 1.3687±0.07 |
| PHIST | Exported protein family 3 | 1.4804±0.09 |
| - | Uncharacterized protein | 1.38715±0.05 |
| H3 | Histone H3 | 1.6045±0.28 |
| FIKK | Serine/threonine protein kinase, FIKK family | 1.3932±0 |
| PM3 | Plasmepsin III | 1.49105±0.17 |
| RESA | Ring-infected erythrocyte surface antigen | 1.28935±0.08 |
| AQP | Aquaglyceroporin | 1.3616±0.03 |
| - | Uncharacterized protein | 1.4352±0.13 |
| - | Uncharacterized protein | 1.31835±0.02 |
| E3 | E3 ubiquitin-protein ligase, putative | 1.31235±0.03 |
| H2B | Histone H2B | 1.77335±0.64 |
| RPS18 | 40S ribosomal protein S18, putative | 1.33095±0.05 |
| UBE2N | Ubiquitin-conjugating enzyme E2 N, putative | 1.29425±0.02 |
| UBE2 | Ubiquitin-conjugating enzyme E2, putative | 1.4351±0.23 |
| UBE2 | Ubiquitin-conjugating enzyme E2, putative | 1.4425±0.24 |
| CDC48 | Cell division cycle protein 48 homologue, putative | 1.25315±0.01 |
| YOP1 | Protein YOP1, putative | 1.41465±0.22 |
| LETM1 | LETM1-like protein, putative | 1.28895±0.06 |
| RPS4 | 40S ribosomal protein S4 | 1.2359±0 |
| EIF3M | Eukaryotic translation initiation factor 3 subunit M | 1.3678±0.19 |
| LAPA | M17 leucyl aminopeptidase | 1.31455±0.11 |
| EF2 | Elongation factor 2 | 1.5163±0.43 |
| VPS35 | Vacuolar protein sorting-associated protein 35 | 1.2781±0.09 |
| - | Uncharacterized protein | 1.3373±0.2 |
| HLYIII | Hemolysin III | 1.29135±0.12 |
| RPL37A | 60S ribosomal protein L37ae, putative | 1.23675±0.06 |
